# Homeostatic regulation of presynaptic NMDA receptor subunit composition modulates action potential driven Ca^2+^ influx into boutons setting the bandwidth for information transfer

**DOI:** 10.1101/2021.09.19.460978

**Authors:** Carla C. Schmidt, Rudi Tong, Nigel J. Emptage

## Abstract

N-Methyl-D-aspartate receptors (NMDARs) play a pivotal role in both short and long-term plasticity. While the functional role of postsynaptic NMDARs is well established, a framework of presynaptic NMDAR (preNMDAR) function is missing. Differences in subunit composition of preNMDARs are documented at central synapses, raising the possibility that subtype composition plays a role in transmission performance. Here, we use electrophysiological recordings at Schaffer collateral - CA1 synapses and Ca^2+^ imaging coupled to focal glutamate uncaging at boutons of CA3 pyramidal neurones to reveal two populations of presynaptic NMDARs that contain either the GluN2A or GluN2B subunit. Activation of the GluN2B population decreases action potential (AP)-evoked Ca^2+^ influx via modulation of small conductance Ca^2+^-activated K^+^ channels (SK-channels) while activation of the GluN2A containing population does the opposite. Moreover, the level of functional expression of each receptor population can be homeostatically modified, bidirectionally affecting short-term facilitation during burst firing, thus providing a capacity for a fine adjustment of the presynaptic integration time window and therefore the bandwidth of information transfer.

## Introduction

The N-Methyl-D-aspartate receptor (NMDAR) is well established as playing a critical role in synaptic plasticity and neural computation (Collingridge, 1987; Collingridge and Bliss, 1987; Kleinschmidt, Bear and Singer, 1987; Morris, Davis and Butcher, 1990; Bashir *et al*., 1991; Malenka and Nicoll, 1993; Regehr and Abbott, 2004; Rebola, Srikumar and Mulle, 2010) with its function extensively explored at the postsynaptic terminal (Tovar and Westbrook, 2002; Wenthold *et al*., 2003; Nakazawa *et al*., 2004; Zhang, Peterson and Liu, 2013; Reese and Kavalali, 2016; Iacobucci and Popescu, 2017; Scheefhals and MacGillavry, 2018). However, although the presence of NMDARs at presynaptic terminals is well documented across a variety of brain areas (Rodríguez-Moreno and Paulsen, 2008; Corlew *et al*., 2008; McGuinness *et al*., 2010; Buchanan *et al*., 2012; Kunz, Roberts and Philpot, 2013; Park, Popescu and Poo, 2014; Larsen *et al*., 2014; Bouvier *et al*., 2015; Gill *et al*., 2015; Banerjee *et al*., 2016; Abrahamsson *et al*., 2017; Padamsey, Tong and Emptage, 2017; Bouvier *et al*., 2018), less is known about the roles they play.

NMDARs are diheteromers comprised of two GluN1 subunits and two GluN2 (GluN2A, GluN2B, GluN2C and GluN2D) or GluN3 (GluN3A and GluN3B) subunits (Paoletti, 2011; Paoletti, Bellone and Zhou, 2013) with GluN2A and GluN2B being the predominant subunits found in the mammalian forebrain (Cull-Candy, Brickley and Farrant, 2001; Yashiro and Philpot, 2008; Paoletti, 2011; Paoletti, Bellone and Zhou, 2013). NMDAR subunit composition creates functional diversity through subunit-specific differences in ion permeability, channel gating and conductance, and coupling to accessory regulatory proteins (Vicini *et al*., 1998; Gielen *et al*., 2009; Rauner and Köhr, 2011; Paoletti, Bellone and Zhou, 2013; Sanz-Clemente, Nicoll and Roche, 2013). At the postsynaptic terminal, NMDAR subunit composition has been linked to distinct processing pathways, often serving opposing roles. For example, GluN2 subunits within hippocampal neurones selectively mediate the direction of plasticity through the regulation of Ca^2+^ influx to engage opposing downstream signalling networks (Fox *et al*., 2006; Shipton and Paulsen, 2014). The inhibition of NMDARs containing the GluN2B subunit prevents long-term depression (LTD) whereas blockade of GluN2A-containing NMDARs inhibits long-term potentiation (LTP) (Liu *et al*., 2004; Massey *et al*., 2004; Bartlett *et al*., 2007; Morishita *et al*., 2007; Yashiro and Philpot, 2008). NMDAR deactivation rate is also known to be dependent on the GluN2 subunit, directly affecting the decay time course of EPSCs and thus the integration properties of the synapse. It is therefore clear that NMDAR subunit composition plays a pivotal role in the tuning of synaptic responses (Monyer *et al*., 1992, 1994; Vicini *et al*., 1998; Popescu and Auerbach, 2003; Lozovaya *et al*., 2004).

Far less well explored is the role of NMDAR subunit composition at the presynaptic terminal, though their importance in various forms of short- and long-term synaptic plasticity is established (Zucker and Regehr, 2002; Sjöström, Turrigiano and Nelson, 2003; McGuinness *et al*., 2010; Bouvier *et al*., 2015; Banerjee *et al*., 2016; Abrahamsson *et al*., 2017; Dore *et al*., 2017; Padamsey, Tong and Emptage, 2017; Lituma *et al*., 2021). For example, presynaptic NMDARs (preNMDARs) are crucial for the induction of LTD at both Schaffer collateral CA1 synapses (Padamsey, Tong and Emptage, 2017) and in the cerebellum (Bidoret *et al*., 2009). Previous efforts to investigate the effects of preNMDAR subunit compositions suggest that preNMDARs containing the GluN2B or GluN2C/D subunits at hippocampal CA3 – CA1 synapses can enhance glutamate release (Prius-Mengual *et al*., 2019), though no mechanistic details are provided. Similarly, in cortical neurones, the GluN2B subunit exhibits a tonic facilitatory effect on spontaneous glutamate release (Chamberlain, Yang and Jones, 2008).

In this study using electrophysiological recordings at Schaffer collateral - CA1 synapses and Ca^2+^ imaging coupled to focal glutamate uncaging at boutons of CA3 pyramidal neurones, we identified two distinct presynaptic NMDAR populations that contain either GluN2A or GluN2B subunits. The acute activation of these preNMDARs either increased (GluN2A) or decreased (GluN2B) action potential (AP)-evoked Ca^2+^ influx via regulation of small conductance Ca^2+^-activated K^+^ channels (SK-channels). Thus, the performance of the synapse depends on the balance of activation of these populations. Further, we show that the preNMDAR populations are plastic and can homeostatically adapt to changes in network activity. The resultant change impacts on short-term facilitation during bursts of action potentials thereby allowing for fine adjustment of the presynaptic integration time window and thus the bandwidth of information transfer.

## Results

### Two distinct populations of preNMDARs bidirectionally modulate AP-evoked Ca^2+^ influx

We examined the action of preNMDARs at the presynaptic terminal of the Schaffer collateral - CA1 synapse in the hippocampus. First, to observe the activation of preNMDARs, we bolus-loaded CA3 pyramidal neurones in hippocampal slices with the Ca^2+^ indicator OGB-1, identified their axonal arbours, and located superficial presynaptic terminals as visually distinct varicosities (Fig. 1A, see Methods). We then used photolytic release of glutamate, supplied by local perfusion of MNI-glutamate, to activate glutamate receptors specifically at the bouton. To prevent the activation of postsynaptic NMDARs, we blocked the activation of AMPA receptors (10 µM NBQX), which are required for the relief of the Mg^2+^ block. The concentration of uncaged glutamate was titrated prior to each experiment at spines located at a similar depth to match the Ca^2+^ transients observed during endogenous evoked release of glutamate (Fig. S1A, see Methods). Similar to the study of Carter and Jahr (2016) in cortical neurones, glutamate photolysis did not cause a significant rise in Ca^2+^, even in conditions of low Mg^2+^ (Fig. S1B). We therefore considered whether Ca^2+^ entry may require additional membrane depolarization. To explore this, we paired glutamate photolysis with an AP, evoked by current injection via the pipette into the neurone under study. To our surprise, pairing an AP with glutamate release between 0.5-5 ms following the onset of the AP, resulted in a decrease in the peak AP-evoked Ca^2+^ transient (APCaT, N = 20 boutons, ΔAPCaT = -0.347 ± 0.049 ΔF/F; Fig. 1B,C, Fig. S1). This decrease was consistent over multiple trials (Fig. 1Biii) but could be completely abolished by bath application of AP5 (50 µM; N = 6 boutons, ΔAPCaT = -0.047 ± 0.024 ΔF/F; Fig. 1C), suggesting the involvement of preNMDARs. Inhibition of metabotropic glutamate receptors (mGluRs) or GABA receptors did not affect the reduction in AP-evoked Ca^2+^ influx (Fig. S2). We further confirmed the presynaptic nature of NMDAR activation by loading the cell with MK-801 (“iMK-801”, 1 mM) through the patch pipette to specifically block preNMDARs within the cell under investigation. This also significantly blocked the decrease in AP-evoked Ca^2+^ transients (N = 12 boutons, ΔAPCaT = -0.137 ± 0.043 ΔF/F; Fig. 1E).

**Figure 1:**
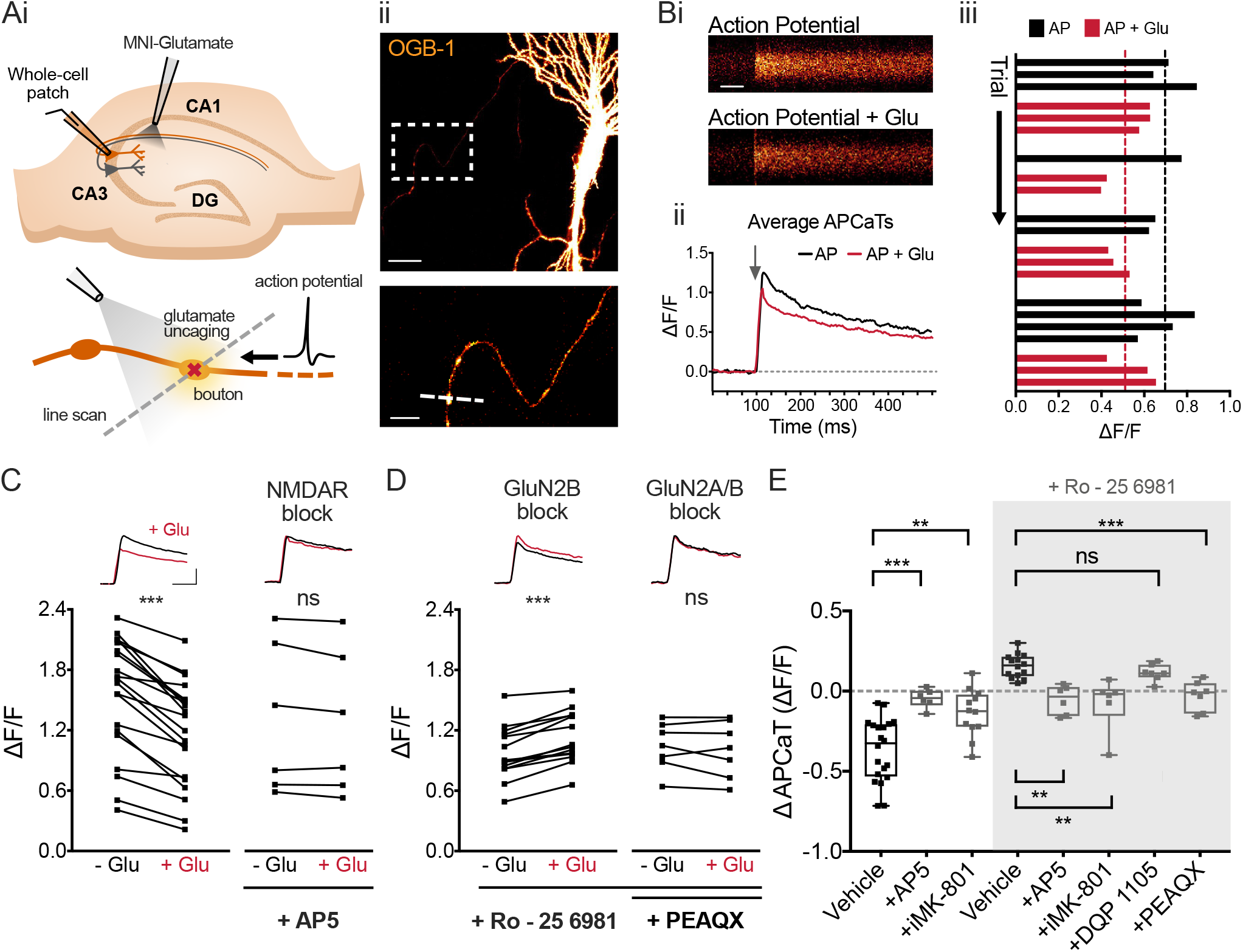
Two distinct populations of preNMDARs bidirectionally modulate AP-evoked Ca^2+^ influx. (Ai) Experimental setup: CA3 cells were patched in whole-cell mode to elicit action potentials via current injection. A glass pipette attached to a picospritzer enabled delivery of MNI-glutamate to the bouton. A 405 nm laser was used for focal uncaging at single boutons. (Aii) *Top*, Example CA3 neurone loaded with OGB-1. The white box indicates presynaptic boutons along the Schaffer collaterals. Scale bar = 20 µm. *Bottom*, Enlarged image of the axon with several boutons. The dashed white line indicates the line scan through the bouton. Scale bar = 5 µm. (Bi) Line scans of AP - evoked Ca^2+^ transients (APCaTs) with or without glutamate photolysis. Scale bar = 50 ms. (Bii) Average APCaTs. Without glutamate = black, With glutamate = red. (Biii) Trial-by-trial peak APCaTs. (C,D) Average peak APCaTs in aCSF, N = 20 (C, left) or in the presence of 50 µM AP5, N = 6 (C, right), 1 µM Ro – 25 6981, N = 14 (D, left) or 1 µM Ro – 25 6981 with 100 nM PEAQX, N = 7 (D, right) to block preNMDARs and specific preNMDAR subunits. PreNMDAR and GluN2B block diminished the decrease in APCaTs. GluN2B block showed an increase in APCaTs. PEAQX blocked the increase in APCaTs. (E) The difference in the peak amplitude between trials with and without glutamate (ΔAPCaT) is shown for control experiments, experiments with 50 µM AP5 or 1 mM intracellular MK-801 (“iMK-801”). Experiments in which GluN2B was blocked with Ro – 25 6981 are highlighted in the grey box (AP5 vs. Vehicle p < 0.001, N = 6, MK-801 vs. Vehicle p = 0.008, N = 12, AP5 vs. Vehicle_Ro_ p < 0.003, N = 6, MK-801 vs. Vehicle_Ro_ p = 0.005, N = 6, PEAQX vs. Vehicle_Ro_ p = 0.006, N = 7, DQP 1105 vs. Vehicle_Ro_ p > 0.99, N = 7; Kruskal-Wallis with *post hoc* Dunn’s test). Error bars represent SEM.

The lack of direct Ca^2+^ entry following glutamate photolysis and the glutamate-induced decrease in APCaTs, are outcomes quite different to those seen for postsynaptic NMDARs. We hypothesised that this may reflect the subunit composition of the preNMDARs. We therefore repeated the experiment with NMDAR antagonists that have a preferential affinity for specific subunits. Bath application of the GluN2B subunit antagonist Ro - 25 6981 (1 µM) produced an increase in APCaTs (N = 12 boutons, ΔAPCaT = 0.157 ± 0.02 ΔF/F; Fig. 1D,E) that was prevented by bath application of AP5 (50 µM), or intracellular MK-801 (1 mM; Fig. 1E). This indicates the presence of a population of preNMDARs that do not contain the GluN2B subunit. Inhibition of the GluN2A subunit using PEAQX (100 nM) completely abolished the increase (N = 7 boutons, ΔAPCaT = -0.029 ± 0.034 ΔF/F; Fig. 1D,E), whereas inhibition of the GluN2C/D subunits using DQP 1105 (50 µM) had no effect (N = 7 boutons, ΔAPCaT = 0.116 ± 0.02 ΔF/F; Fig. 1E). PEAQX application alone preserved the decrease in APCaTs (Fig. S2), however, to a more modest extent than in control conditions, most likely due to the lower specificity and partial inhibition of the GluN2B subunit (Auberson *et al*., 2002; Feng *et al*., 2004).

These data suggest that two populations of NMDARs are present at CA3 presynaptic terminals, with one that contains the GluN2A subunit and the other the GluN2B subunit.

### PreNMDARs modulate AP-evoked Ca^2+^ influx by the activation of SK-channels

We next explored how the subunit composition of preNMDARs was able to differentially influence the AP-evoked Ca^2+^ influx. At the postsynaptic terminal small-conductance Ca^2+^-activated K^+^ channels (SK-channels) are known to form a Ca^2+^-mediated negative feedback loop with NMDARs to reduce Ca^2+^ influx into synaptic terminals (Ngo-Anh *et al*., 2005; Faber, 2010; Griffith, Tsaneva-Atanasova and Mellor, 2016). Moreover, SK-channels are well- established as shaping the action potential wave form (Xia *et al*., 1998; Sah and Faber, 2002; Faber and Sah, 2003, 2007; Bond, Maylie and Adelman, 2005), impacting on the dynamics of presynaptic voltage-gated Ca^2+^ channels (VGCCs) and ultimately release probability (P^r^) (Bean, 2007; Hoppa *et al*., 2014; Chéreau *et al*., 2017). Thus, we hypothesised that NMDAR/SK-channel pathways may also be present in CA3 presynaptic terminals and that these are able to modulate the action potential evoked Ca^2+^ influx.

Bath application of the selective SK-channel blocker Apamin (1 µM) abolished both the glutamate photolysis-induced decrease (N = 13 boutons, ΔAPCaT = -0.097 ± 0.037 ΔF/F; Fig. 2A,D) and increase (N = 9 boutons, ΔAPCaT = -0.046 ± 0.035 ΔF/F; Fig. 2B,D), observed following the application of Ro – 25 6981, suggesting that preNMDARs of different subunit composition converge upon a common pathway. We also thought it important to assess whether intracellular stores formed part of the pathway and so applied CPA (15 µM) and ryanodine (20 µM), however these did not affect the increase in APCaTs (N = 8 boutons, ΔAPCaT = 0.144 ± 0.029 ΔF/F; Fig. 2C,D).

**Figure 2:**
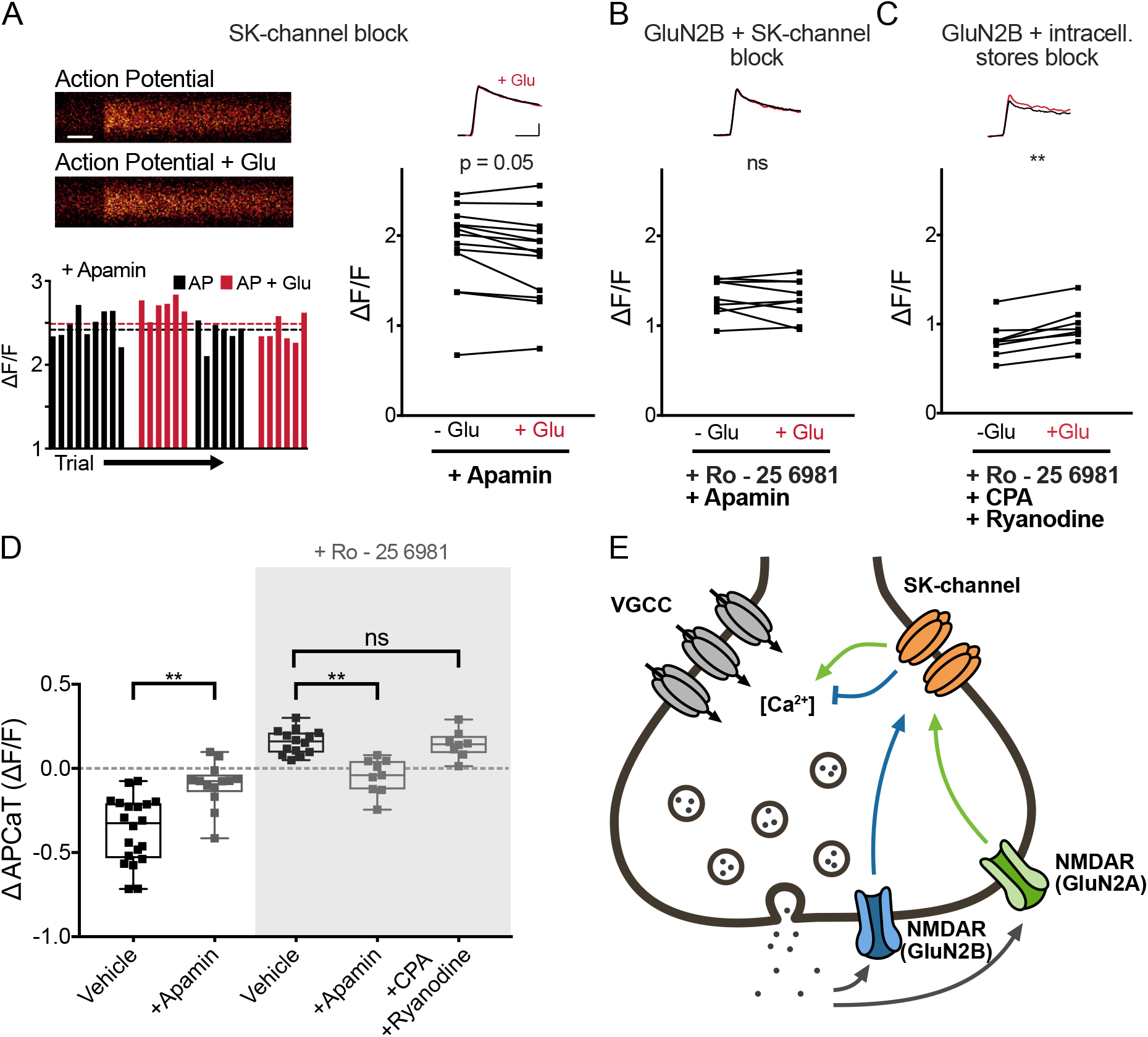
PreNMDARs modulate AP-evoked Ca^2+^ influx via the activation of SK-channels. (A) *Top left*, Line scans of APCaTs following SK – channel inhibition with 1 μM Apamin. Scale bar = 50 ms. *Bottom left*, Trial-by-trial peak APCaTs. Without glutamate = black, With glutamate = red. *Right*, Average peak APCaTs in 1 µM Apamin. The decrease in peak APCaTs was diminished after SK - channel inhibition, N = 13. (B,C) Similarly, the Ro – 25 6981 dependent increase in peak APCaTs was abolished after application of 1 µM Apamin, N = 9. Inhibition of intracellular Ca^2+^ stores using CPA (15 µM) and ryanodine (20 µM) did not affect the increase, N = 8. (D) The difference in the peak amplitude of APCaTs is shown for experiments in aCSF, N = 20 (from Fig. 1) and experiments with 1 µM Apamin. Experiments in which GluN2B was blocked with Ro – 25 6981 are highlighted in the grey box (Apamin vs. Vehicle p < 0.001, Apamin vs. Vehicle_Ro_ p = 0.001, CPA & Ryanodine vs. Vehicle_Ro_ p > 0.99; Kruskal-Wallis with *post hoc* Dunn’s test). Error bars represent SEM. (E) Proposed molecular mechanism for preNMDAR and SK-channel mediated boutonal Ca^2+^ dynamics. The release of glutamate activates two different preNMDAR populations, containing either GluN2A or GluN2B, which causes local Ca^2+^ influx and leads to the modulation of SK-channels, resulting in an increase or a decrease in Ca^2+^ influx, respectively, likely through voltage-gated Ca^2+^-channels (VGCCs).

We conclude from these observations that following the release of glutamate, two populations of preNMDARs become activated, each triggering a cascade of intracellular signalling events which converge upon SK-channels (Fig. 2E). The SK-channels influence the duration of the AP and consequently the opening of VGCCs. Whether an increase or a decrease in the AP-evoked Ca^2+^ influx occurs reflects the dominance of one pathway over the other.

### PreNMDAR modulation of AP-evoked Ca^2+^ influx results in use-dependent modulation of short-term facilitation

Changes in presynaptic Ca^2+^ signalling predominantly impact on neurotransmitter release and short-term plasticity (STP). In particular, the temporal profile of the Ca^2+^ concentration within the bouton is linked to the expression of short-term facilitation (Zucker and Regehr, 2002). We therefore hypothesised that preNMDAR-mediated modulation of Ca^2+^ influx acts to modify short-term facilitation. Furthermore, the requirement of glutamate for the activation of the preNMDARs ensures that modulation is use-dependent, i.e. conditioned by the recent history of release events, rather than simply the presence of an AP train alone.

Figure 3A illustrates potential outcomes of use-dependent modulation of short-term facilitation. The activation of the GluN2A dominant pathway forms a positive feedback loop in which the release of glutamate will augment neurotransmitter release during a train of APs. This mechanism ensures that multiple release events occur for an AP burst, increasing the robustness of information transmission, by extending the impact of the burst in time, thereby increasing the integration time window at the postsynaptic neurone. By contrast, the GluN2B dominant pathway forms a negative feedback loop that reduces the increase in P_r_ during bursts of APs. For this pathway to be active, neurotransmitter release must of course first occur, a condition that ensures that some neurotransmitter is released. This use-dependent ‘clamping’ of short-term facilitation prevents excessive vesicle depletion (i.e. short-term depression) and resets the synapse for subsequent AP trains. This will enhance information transfer within short integration time windows. The precise balance between GluN2A and GluN2B subpopulations can therefore set the optimal transmission mode for a given synapse.

**Figure 3:**
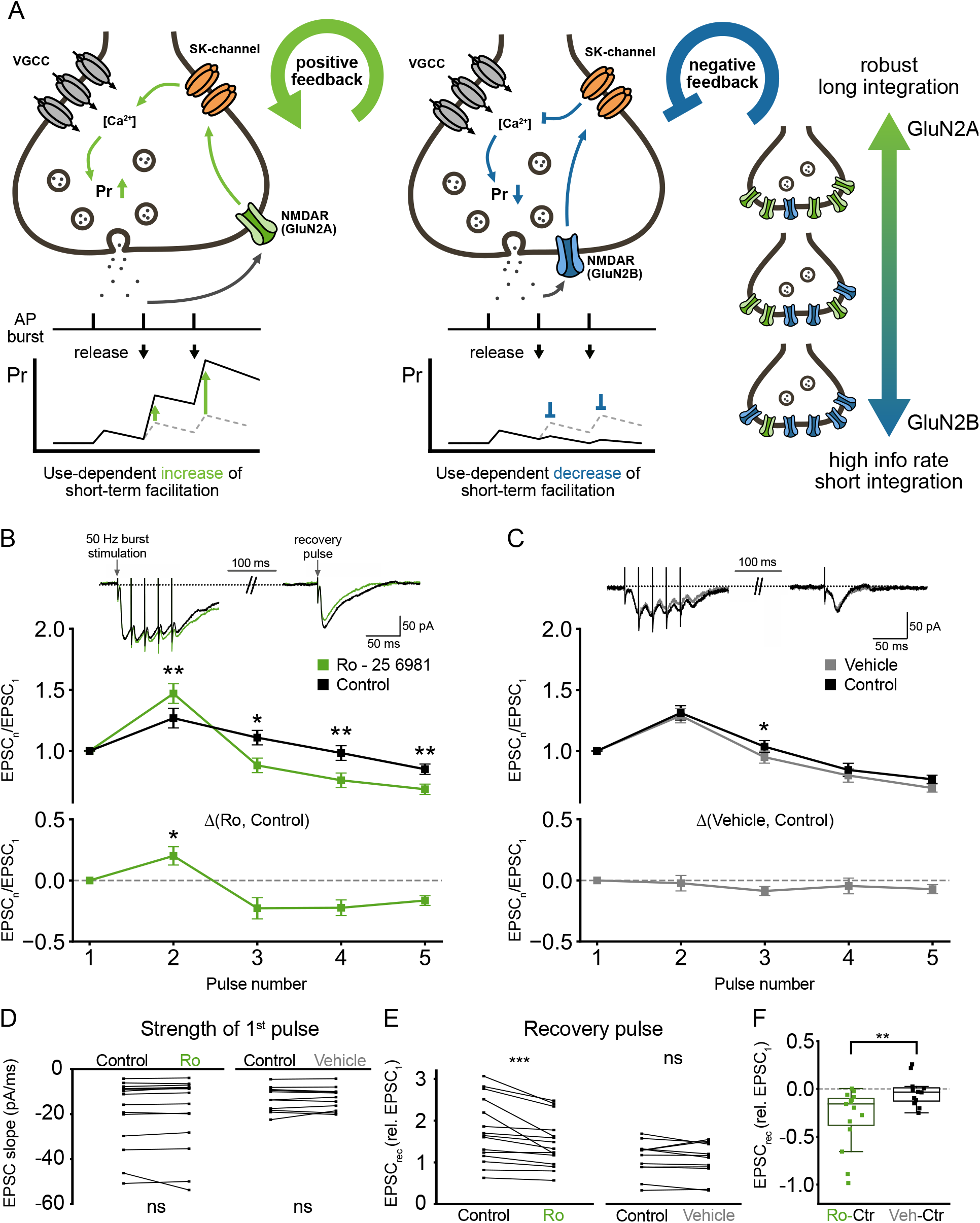
PreNMDAR modulation of AP-evoked Ca^2+^ influx results in use-dependent modulation of short-term facilitation. (A) Potential consequence of use-dependent augmentation of short-term facilitation. The activation of the GluN2A dominant pathway leads to a use-dependent increase in short-term facilitation which ensures that multiple quanta of release will be transmitted by a given AP burst. This results in an extension of the AP burst in time, which increases the integration time window at the postsynaptic neurone. The GluN2B dominant pathway, following glutamate release, acts to prevent the increase in P_r_ caused by short-term facilitation. This use-dependent ‘clamp’ of short-term facilitation prevents excessive vesicle depletion (i.e. short-term depression) and resets the synapse for subsequent AP trains allowing for fast information transfer with short integration time windows. (B) *Top*, Sample traces of burst stimulation (5 APs at 50 Hz) before (black) and after the addition of 1 μM Ro – 25 6981 (green) to block GluN2B containing preNMDARs. *Middle*, Average normalised slope of the EPSC during burst stimulation before and after the addition of Ro – 25 6981. Application of Ro – 25 6981 significantly increased the magnitude of short-term facilitation (N = 15 cells; p = 0.001 for pulse 2; Wilcoxon signed rank test) and enhanced the magnitude of short-term depression (N = 15 cells; p < 0.05 for pulse 3-5; Wilcoxon signed rank test). *Bottom*, The difference in the normalised EPSC slope of pulse 1-5 before and after the addition of Ro - 25 6981 (significance indicates test against Vehicle in (C); p < 0.05 for pulse 2; Mann-Whitney test. (C) Same as B for Vehicle (aCSF alone, N = 11 cells). (D) The initial EPSC slope does not change during the experiment (N = 15 cells (control), p = 0.08; N = 11 cells (Vehicle), p > 0.99; Wilcoxon signed rank test). (E,F) The magnitude of the recovery pulse given 100 ms after the burst was significantly decreased when GluN2B activity was inhibited with Ro - 25 6981 but not in control experiments (Ro - 25 6981: p < 0.001; Vehicle: p = 0.34; Wilcoxon signed rank test; Ro vs vehicle: p < 0.01, Mann Whitney test).

To investigate whether preNMDAR subunit populations differentially regulate short-term facilitation, we recorded from CA1 neurones in response to bursts of APs (5 pulses at 50 Hz) elicited at the Schaffer collateral inputs. We also included a single recovery pulse 100 ms after the burst (Fig 3B, top) to examine the speed of recovery from short-term depression. 1 mM MK-801 was included in the patch electrode to block postsynaptic NMDARs.

AP trains elicited short-term facilitation, followed by a slow decline (Fig. 3B,C) as previously reported for these synapses (Stevens and Wang, 1995; Dobrunz, Huang and Stevens, 1997; James *et al*., 2006; Sun and Dobrunz, 2006; Bartley and Dobrunz, 2015). Application of the GluN2B subunit blocker Ro - 25 6981 (1 µM) significantly increased short-term facilitation of the second pulse and enhanced short-term depression of subsequent pulses (Fig. 3B). Short-term plasticity remained unchanged when aCSF alone was perfused (Fig. 3C). Basal P_r_ is a key determinant of the short-term behaviour of a synapse. We therefore wished to rule out the possibility that application of Ro - 25 6981 caused a change in P_r_. For this we compared the magnitude of the first pulse in each burst (Ro - 25 6981: N = 15 cells, p = 0.09; Vehicle: N = 11 cells, p > 0.99; Wilcoxon signed rank test; Fig. 3D). We did not find a significant difference. We next examined the recovery from short-term depression. The magnitude of the single recovery pulse given 100 ms after the burst was significantly decreased when GluN2B activity was inhibited with Ro - 25 6981 (Ro - 25 6981: N = 15 cells, p < 0.001; Vehicle: N = 11 cells, p = 0.34; Wilcoxon signed rank test; Fig. 3E), but not in control vehicle experiments (Ro - 25 6981 vs. Vehicle: p = 0.01; Mann-Whitney U test; Fig. 3F).

Consistent with our hypothesis the results show that preNMDARs containing different subunits differentially regulate short-term facilitation during bursts of APs, a mechanism that allows for fine adjustment of the information transfer properties of the terminal.

### Network activity shifts the balance between the GluN2A and GluN2B subpopulations

What determines the contribution of each preNMDAR subpopulation? Given the functional role outlined above, the balance between subpopulations should ideally be adjustable with respect to network activity, with sparse activity favouring the robust transmission mode implemented by the GluN2A subunit and increased activity favouring the GluN2B subunit. We tested this by globally increasing (1 µM gabazine) or decreasing (10 µM NBQX and 50 µM AP5) network activity for 48-72 hours (Fig. 4A), an established protocol for induction of homeostatic plasticity (Turrigiano, 2012). We then measured the modulation of APCaTs.

**Figure 4:**
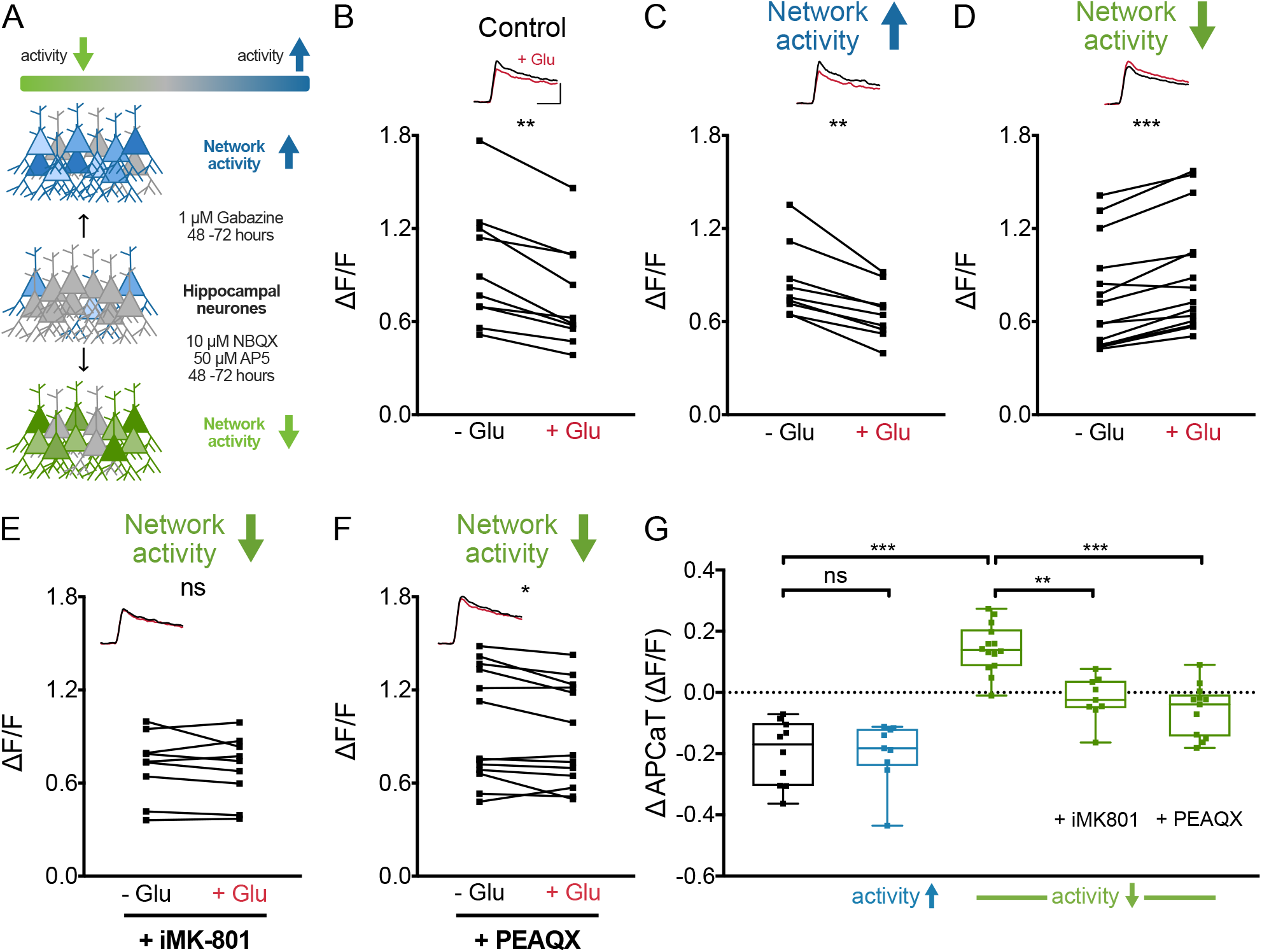
Network activity shifts the balance between GluN2A and GluN2B subpopulations. (A) Schematic of the experimental conditions. Network activity was either increased or decreased via application of 1 µM Gabazine or 10 µM NBQX and 50 µM AP5 for 48-72 hours, respectively. (B-D) Peak APCaTs in control conditions, N = 10 (B) and after increasing, N = 9 (C) or decreasing, N = 14 (D) network activity. Decreasing network activity led to an increase in peak APCaTs (D). This increase was abolished with intracellular MK-801, N = 9 and PEAQX, N = 13 (E,F). (G) Summary of the regulation of subpopulation balance by network activity (Vehicle_lowactivity_ vs. Control p < 0.001, Vehicle_lowactivity_ vs. iMK-801 p = 0.002, PEAQX vs. Vehicle_lowactivity_ p < 0.001; Kruskal-Wallis with *post hoc* Dunn’s test). Error bars represent SEM.

Increased network activity resulted in an overall decrease in APCaTs (N = 9 boutons, ΔAPCaT = -0.2 ± 0.034 ΔF/F; Fig. 4C) which was not significantly different from control experiments (Gabazine vs. CTR p > 0.99; Kruskal-Wallis with *post hoc* Dunn’s test; Fig. 4B,C,G). In contrast, decreasing network activity resulted in a substantial increase in APCaTs (N = 14 boutons, ΔAPCaT = 0.144 ± 0.021 ΔF/F; Fig. 4D) following photolysis of glutamate (AP5/NBQX vs. CTR p < 0.001; Kruskal-Wallis with *post hoc* Dunn’s test; Fig. 4G). We verified that this increase is caused by an increase in GluN2A containing preNMDARs via intracellular MK-801 (N = 9 boutons, ΔAPCaT = -0.019 ± 0.024 ΔF/F; Fig. 4E,G) and application of PEAQX (N = 13 boutons, ΔAPCaT = -0.057 ± 0.023 ΔF/F; Fig. 4F,G). Application of PEAQX did not unmask a decrease in APCaTs (Fig. 4G), suggesting that the extensive silencing of network activity has caused a shift in the dominant subpopulation expression such that far fewer GluN2B containing receptors remained.

## Discussion

Here we show that the activation of two preNMDAR populations, each with a distinct subunit composition, bidirectionally regulate Ca^2+^ dynamics and STP at Schaffer collateral boutons. Activation of preNMDARs containing the GluN2B subunit form a negative feedback-loop, via SK-channels, that decrease the Ca^2+^ influx that occurs during an action potential leading to a reduction of short-term facilitation during high frequency firing. Activation of preNMDARs containing the GluN2A subunit increase the action potential driven Ca^2+^ influx into the bouton, which results in the reinforcement of short-term facilitation during burst firing.

We show that the GluN2B pathway is predominant in conditions of high network activity, as commonly observed in hippocampal slice preparations (De Simoni, Griesinger and Edwards, 2003), whereas GluN2A signalling dominates when network activity is low. This may explain why detection of glutamate evoked Ca^2+^ influx through preNMDARs has proven challenging (Christie and Jahr, 2008, 2009; Carter and Jahr, 2016) including in our own study (Fig. S1B). However, when GluN2B subunits were blocked, we were able to observe a small, but significant increase in Ca^2+^, even in the absence of APs (Fig. S3). Hence, our data support the idea that synaptic terminals flexibly adjust the balance between GluN2A and GluN2B subpopulations depending on the activity state of the network. Whether this balance depends additionally on local activity, meaning the unique pre- and postsynaptic activity experienced by each individual synapse, remains to be explored, though synapse-specific differences in the expression of preNMDARs within the same neurone have been reported in cortex (Buchanan *et al*., 2012). This would endow the presynaptic terminal with exceptional computational flexibility and adds support to a recently proposed model for presynaptic computation (Tong, Emptage and Padamsey, 2020). Whether the upregulation of GluN2A containing preNMDARs during phases of low network activity results from local translation (Holt and Schuman, 2013; Holt, Martin and Schuman, 2019) or membrane trafficking (Aoki *et al*., 2003; Barria, 2003; Lau and Zukin, 2007) also remains to be investigated.

Within the hippocampus a large variety of NMDAR subunits and isoforms can be found, though a receptor’s functional performance is thought to be mostly determined by the GluN2 subunit (Gielen *et al*., 2009; Paoletti, 2011; Paoletti, Bellone and Zhou, 2013; Sanz-Clemente, Nicoll and Roche, 2013). In our study, we focused on GluN2A and GluN2B as they are the predominantly expressed isoforms in the adult hippocampus. In contrast, GluN2C and GluN2D expression levels in the adult brain are considerably lower and most prominent in the cerebellum and the brainstem (Monyer *et al*., 1994; Paoletti, 2011; Paoletti, Bellone and Zhou, 2013; Sanz-Clemente, Nicoll and Roche, 2013; Wyllie, Livesey and Hardingham, 2013) or astrocytes (Chipman *et al*., 2021). Although most studies have focused on diheteromeric GluN1/GluN2 receptors, the importance of triheteromeric NMDARs is also recognised (Cull-candy and Leszkiewicz, 2004; Köhr, 2006; Paoletti and Neyton, 2007; Rauner and Köhr, 2011; Dubois *et al*., 2016; Stroebel, Casado and Paoletti, 2018), raising the possibility for even broader functional diversity. Here, our observations for the preNMDAR subpopulation involved in the negative feedback loop (containing GluN2B) are consistent with both a GluN1/GluN2B diheteromer or GluN1/GluN2A/GluN2B triheteromer expression and therefore requires a more detailed pharmacological and genetic dissection of subunit composition.

It is well established that SK-channels are activated by Ca^2+^ influx through NMDARs (Shah and Haylett, 2002; Faber, Delaney and Sah, 2005; Lin *et al*., 2008) which reduces Ca^2+^ influx through VGCCs (Bond, Maylie and Adelman, 2005; Faber and Sah, 2007; Griffith, Tsaneva-Atanasova and Mellor, 2016). The negative feedback between NMDARs and SK channels has been carefully studied at the postsynaptic terminal (Ngo-Anh *et al*., 2005; Faber, 2010; Griffith, Tsaneva-Atanasova and Mellor, 2016), but not at the presynaptic terminal, even though SK-channels and NMDARs are known to be colocalised there (Ngo-Anh *et al*., 2005; Nanou *et al*., 2013). In this study we confirm the presence of the negative feedback interaction at the presynaptic terminal and link it specifically to the GluN2B subunit of the NMDAR. Additionally, we have shown that the GluN2A subunit is required for the formation of a positive feedback loop with SK-channels. This subunit-dependent bidirectional signalling is analogous with the well characterised roles of postsynaptic NMDARs in LTP and LTD (Liu *et al*., 2004; Massey *et al*., 2004; Fox *et al*., 2006; Shipton and Paulsen, 2014).

How do the two preNMDAR populations produce their differential effects upon the SK-channels? While GluN2A and GluN2B show similar characteristics for Ca^2+^ permeability, sensitivity to Mg^2+^ blockade, and channel conductance, GluN2A-containing receptors are coupled to distinct downstream signalling networks (Paoletti, Bellone and Zhou, 2013; Sanz-Clemente, Nicoll and Roche, 2013). SK-channels are part of large protein complexes and are co-assembled with protein kinase CK2 and phosphatase PP2A, which have opposing effects on SK-channel activity (Bildl *et al*., 2004; Allen *et al*., 2007; Luján, Maylie and Adelman, 2009): while CK2 phosphorylates SK-channel bound Calmodulin (CaM) resulting in faster channel deactivation and reduced Ca^2+^ sensitivity, PP2A dephosphorylates CaM causing enhanced Ca^2+^ sensitivity (Luján, Maylie and Adelman, 2009). Therefore, one mechanism could be that the activation of GluN2A receptors increases CK2 activity within the bouton, leading to an inhibition of SK-channels thus prolonging membrane depolarization and consequently Ca^2+^ influx (Ngo-Anh *et al*., 2005; Griffith, Tsaneva-Atanasova and Mellor, 2016), whereas the activation of the GluN2B containing subpopulation engages the PP2A signalling pathway leading to a reduction in the depolarization caused by the AP. Differential activation of kinase and phosphatase pathways lie at the heart of the expression of LTP/LTD at the postsynaptic terminal where competition between kinases and phosphatases is mediated by differences in Ca^2+^ sensitivity and signalling molecules that are coupled to distinct NMDAR isoforms (Bear and Malenka, 1994; Coussens and Teyler, 1996; Yang, Tang and Zucker, 1999; Malenka and Bear, 2004; Grey and Burrell, 2010). While this is an attractive possibility with NMDARs able to activate both CK2 and PP2A, a causal link to the regulation of SK-channels is not shown in this study.

Finally, the differential expression patterns of preNMDAR subunits and the resulting functional heterogeneity we have identified may account for some of the differences in the literature. Functional and anatomical evidence for preNMDARs has been available for a number of years (Pittaluga and Raiteri, 1990; Duguid and Smart, 2004; Janssen *et al*., 2005; McGuinness *et al*., 2010), however, the existence of functional receptors has been challenged as some studies report NMDAR-dependent Ca^2+^ transients in boutons (McGuinness *et al*., 2010; Buchanan *et al*., 2012) while others do not (Christie and Jahr, 2008, 2009; Carter and Jahr, 2016). Here we are only able to detect a preNMDAR-dependent modulation of Ca^2+^ when activation was paired with APs. Furthermore, only after isolating the GluN2A subunit is Ca^2+^ influx through the preNMDARs unmasked (Fig. S3). It therefore seems prudent to review data with careful oversight of preNMDAR subunit composition.

## Acknowledgements

We would like to thank Zahid Padamsey for technical guidance, helpful discussions and feedback. C.C.S. received funding from the University of Oxford and the Medical Research Council UK. R.T. was jointly funded by the Clarendon Fund, University of Oxford and the Medical Research Council UK.

## Author contributions

C.C.S and R.T conducted and analysed experiments. Manuscript and figures were prepared by C.C.S. and R.T. and edited and approved by N.J.E. Experiments were conceptualized and designed by all authors.

## Declaration of interests

The authors declare no competing interests.

## Methods

### Organotypic hippocampal slices

Transverse 350 μm organotypic hippocampal slices were prepared from male Wistar rats (P7-P8). After dissection, slices were cultured on Millicell CM membranes and maintained in culture media at 37°C for 7-14 days prior to use. Culture media was comprised of 78.8% Minimum Essential Media, 20% heat-inactivated horse serum, 30 mM HEPES, 26 mM D-Glu, 5.8 mM NaHCO_3_, 1 mM CaCl_2_, 2 mM MgSO_4_ and 1% B-27. The media was replaced every 2-3 days to ensure optimal and constant conditions for the slices. During the experiments slices were superfused with oxygenated (95% O2/5% CO_2_) artificial cerebrospinal fluid (ACSF; composition in mM: 145 NaCl, 2.5 KCl, 2 MgCl_2_, 3 CaCl_2_, 1.2 NaH2PO_4_, 16 NaH2CO_3_, 11 glucose).

### Pharmacology

200 nM NBQX was added to the ACSF to prevent hyperactivity in organotypic slices. In order to block presynaptic NMDARs, 50 μM AP5 was either washed in or included in the aCSF from the start. Ro - 25 6981 (1 µM), PEAQX (100 nM) and DQP 1105 (50 μM) were used to specifically block NMDAR subunits. For SK-channel inhibition, slices were preincubated in 1 µM Apamin, which was maintained throughout the experiment. During electrophysiology experiments, slices were perfused with Ro – 256981 (1 µM) for at least 10 minutes to ensure reliable block of GluN2B. To manipulate global network activity and to induce homeostatic plasticity, slices were incubated for 48 – 72 hours with either gabazine (1 µM) or NBQX (10 µM) / AP5 (50 µM), before imaging.

### Electrophysiology and Imaging

Electrophysiological data was recorded in whole-cell patch clamp mode with WinWCP (Strathclyde Electrophysiology 851 Software) and analyzed using Clampfit (Axon Instruments). Cells were imaged with a LEICA DMLFSA microscope fitted with a 63x water-immersion objective (HCX APO L 63x/0.9W U-V-I, Leica) and a LEICA TCS SP2 confocal scan head. For Ca^2+^ imaging, superficial CA3 pyramidal cells were patched with low-resistance patch electrodes (4-8 MΩ) containing the Ca^2+^-sensitive dye Oregon Green 488 BAPTA-1 (OGB-1; 0.5-1 mM) for 1-5 min. We waited at least 30-45 min for the indicator to reach diffusional equilibrium in the axon. We closely monitored signal intensity and we did not see an increase in the basal fluorescence intensity of the dye over the course of the experiments. Furthermore, dye saturation in the axon was examined by testing the summation of Ca^2+^ responses of two APs given in quick succession. After successful identification of the axon and superficial boutons, cells were repatched. Subsequently, line scans through boutons were performed and synchronized to intrasomatically stimulated APs triggered by injecting step currents (0.5–2 pA) of 10 ms duration. We recorded 10-15 successive trials per condition. Ca^2+^ responses were averaged within trials and the peak response was extracted. Ca^2+^ transients in boutons were analysed using ImageJ and expressed as the fractional change in fluorescence (ΔF/F):

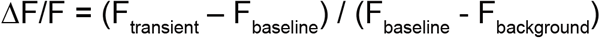

To elicit high-frequency action potential bursts, a monopolar tungsten electrode encased in a glass pipette was positioned in stratum radiatum to stimulate Schaffer collaterals. CA1 pyramidal cells were stimulated with 5 pulses at 50 Hz followed by a single pulse given 100 ms from the end of burst. Patch electrodes contained 1 mM MK-801 to block postsynaptic NMDARs.

### Glutamate Uncaging

Before each experiment, we titrated the intensity of the uncaging laser (405 nm UV-laser) to detect a robust rise in Ca^2+^ (0.5 - 1 ΔF/F) in distal dendritic spines (Fig. S1). This calibration ensures the release of glutamate at physiologically relevant concentrations and minimises phototoxicity. After identification of superficial boutons, a glass pipette (4-8 MΩ) filled with MNI-glutamate (10 mM) and connected to a picospritzer was placed close to the boutons (within 20 µm just above the surface of the slice) to ensure focal delivery of the MNI-glutamate. Since intrasomatically induced APs take time to reach the presynaptic terminals and to release glutamate, we set the glutamate uncaging to occur 0.5-5 ms after the induced action potential (Fig. S1).

### Statistical analysis

ImageJ and Graphpad Prism 7 were used for analysis, graphing and statistical testing. Data was analyzed with two-tailed Mann-Whitney U test, Wilcoxon signed rank test or Kruskal-Wallis with post-hoc Dunn’s test for multiple comparisons. Data is reported as mean ± standard error of the mean (SEM). Significance is denoted as follows: *p < 0.05, **p < 0.01, ***p < 0.001.

---

**Figure S1:**
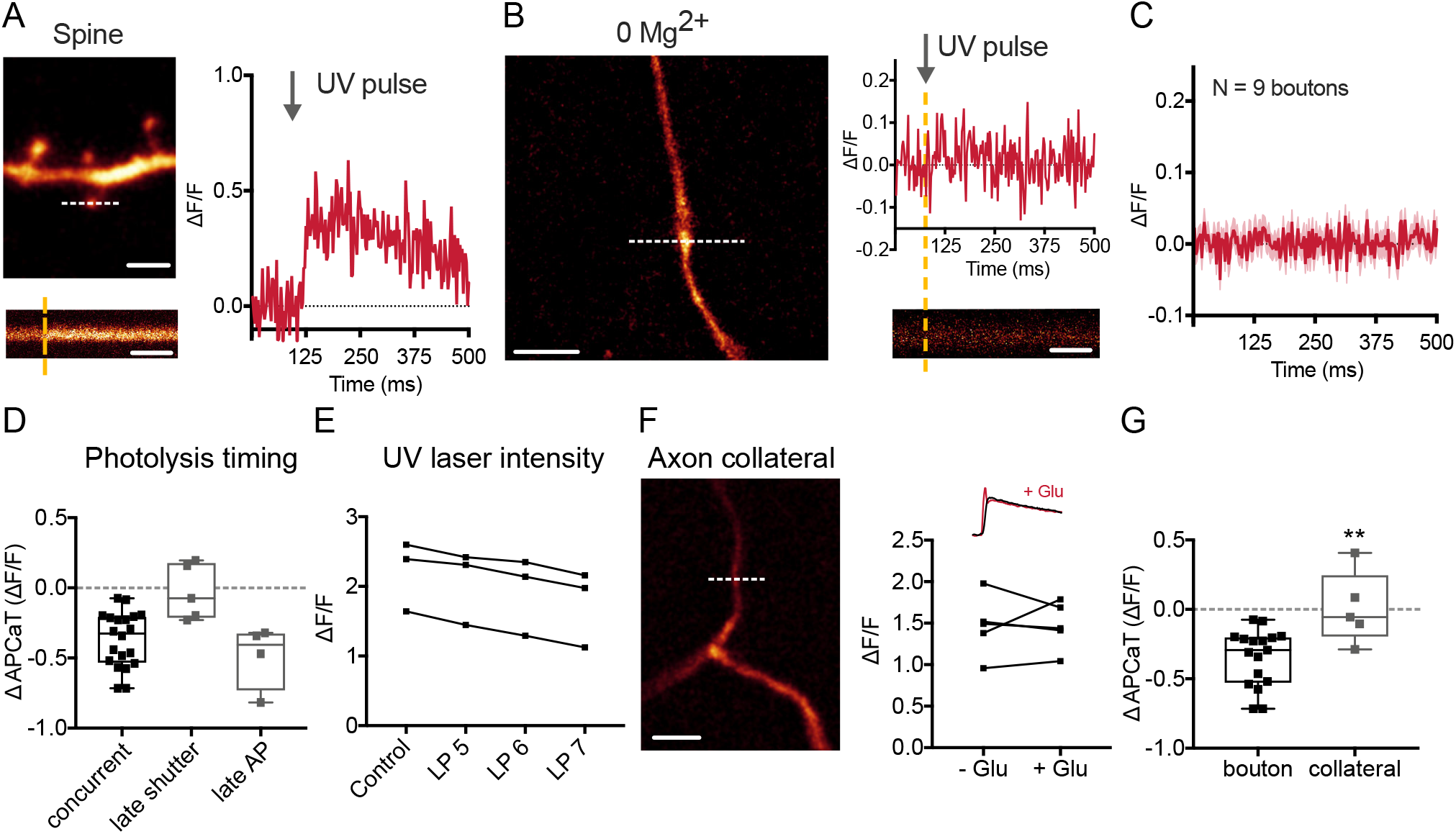
Glutamate uncaging alone does not elicit preNMDAR Ca^2+^ influx and requires concurrent AP firing. Related to Figure 1. (A) *Top*, An example image of a dendrite from a CA3 neurone loaded with OGB-1. Scale bar = 5 µm. *Bottom*, Line scan through a dendritic spine as indicated above. Scale bar = 125 ms. *Right*, The UV uncaging laser was titrated at small spines to elicit a response of approximately 0.5 ΔF/F. (B) *Left*, Example of an axon and bouton. Scale bar = 10 µm. *Right*, Example trace and line scan of the bouton in Bi. Glutamate uncaging in low Mg^2+^ did not elicit a Ca^2+^ response at Schaffer collateral boutons. (C) Average trace from 9 different boutons across three cells. (D) Difference in peak APCaTs between trials with and without glutamate photolysis for concurrent experiments (photolysis 0.5 – 5 ms after AP), N = 5, late shutter (20 ms after AP), N = 5, and late AP (20 ms post photolysis), N = 4. The decrease in average peak APCaTs under control conditions is still observable when the AP was delayed but abolished when the shutter opening was delayed. (E) Increasing UV laser intensity linearly decreased APCaTs, likely due to an increase in the concentration of uncaged glutamate. Laser power (LP) 5 – 7 in arbitrary units, N = 3. (F,G) Photolysis experiments were performed at axon collaterals instead of boutons, N = 5. Scale bar = 10 µm. No change in AP-evoked Ca^2+^ transients could be detected following glutamate photolysis at the collateral (Bouton vs. Collateral p = 0.004; Mann Whitney test). Error bars represent SEM.

**Figure S2:**
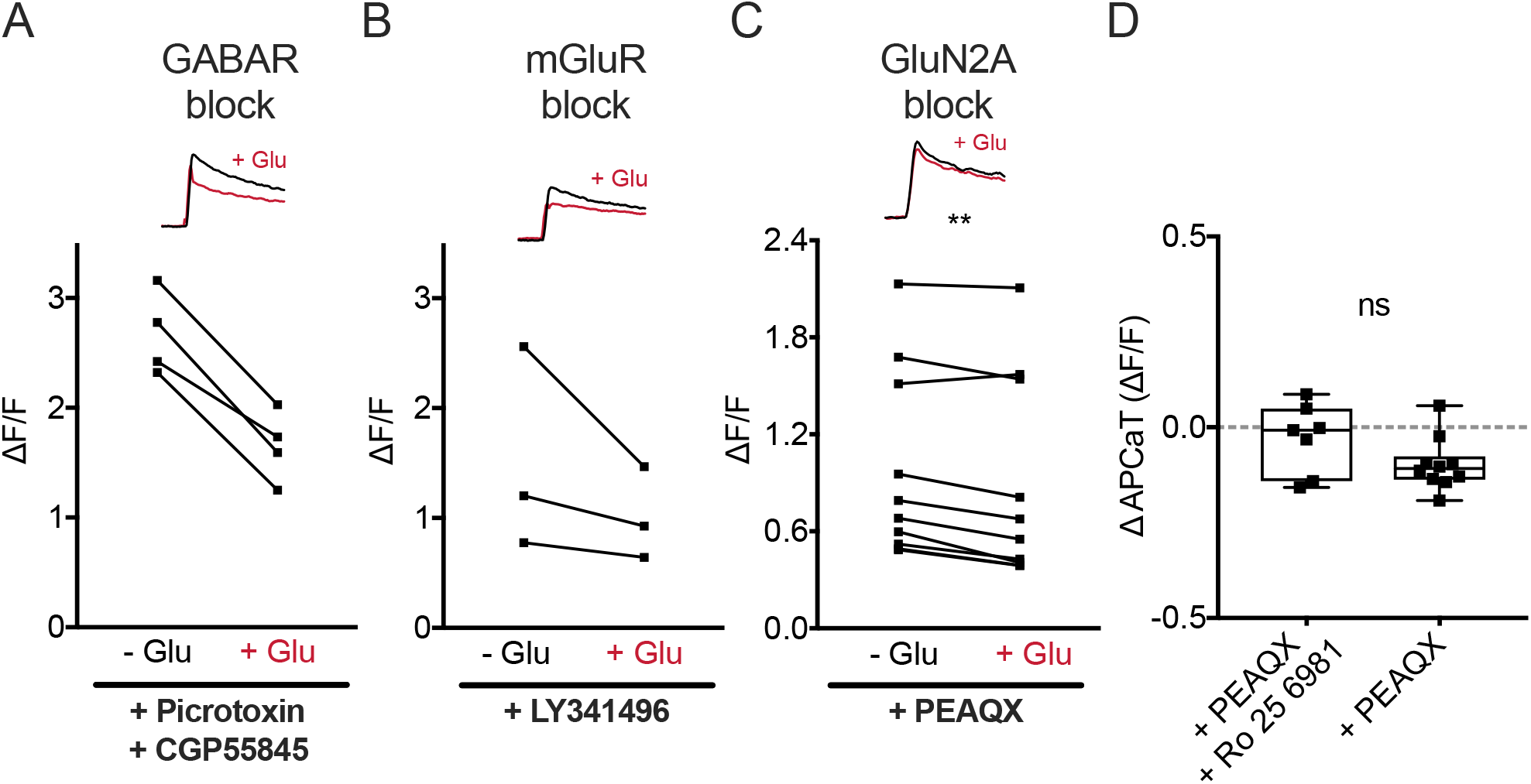
Modulation of AP-evoked Ca^2+^ influx is independent of GABA receptors and mGluRs. Related to Figure 1. (A,B) Inhibition of GABA receptors, N = 4 or metabotrobic glutamate receptors (mGluRs), N = 3, with Picrotoxin (30 µM), CGP5845 (5 µM) and LY341495 (100 µM) did not block the decrease in APCaTs. (C,D) PEAQX (100 nM) application preserved the decrease in Ca^2+^ signal, N = 10. Error bars represent SEM.

**Figure S3:**
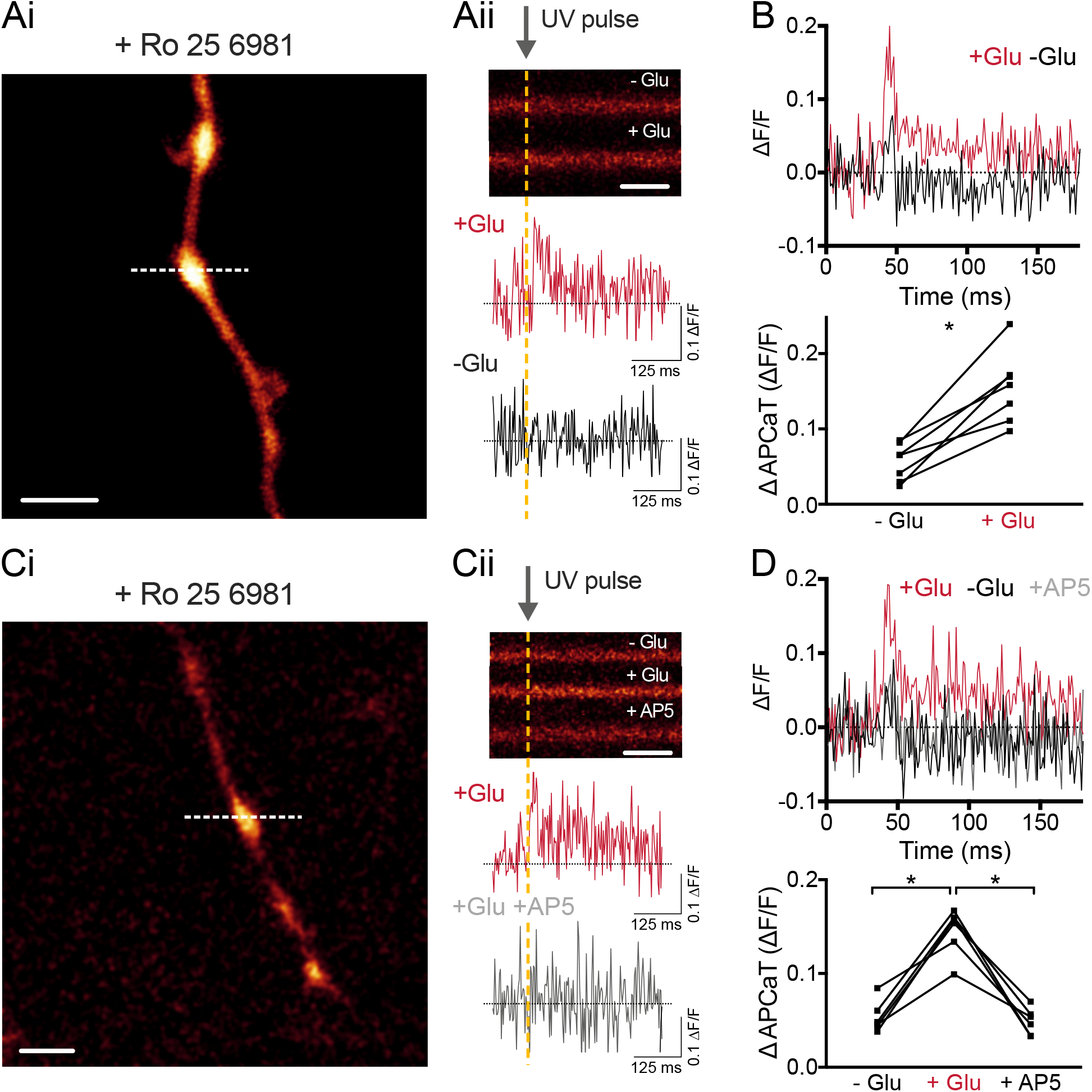
Glutamate uncaging induced Ca^2+^ influx at Schaffer collateral boutons following GluN2B inhibition. Related to Figure 1. (Ai) Line scans through boutons of CA3 neurones loaded with OGB-1. Scale bar = 5 µm. (Aii) Blockage of GluN2B subunits with 1 μM Ro – 25 6981 unmasked an increase in Ca^2+^ following glutamate uncaging in low Mg^2+^. Scale bar = 50 ms. (B) *Top*, Average Ca^2+^ transient following glutamate uncaging (N = 7 boutons, p = 0.016; Wilcoxon signed rank test). *Bottom*, GluN2B inhibition unmasked an increase in Ca^2+^ influx at all boutons tested. (C,D) Application of AP5 blocked the Ca^2+^ increase (N = 6 boutons, -Glu vs +Glu: p = 0.03, +Glu vs AP5: p = 0.03; Wilcoxon signed rank test).

## Notes

### Competing Interest Statement

The authors have declared no competing interest.

